# Inhibiting IFT dynein with ciliobrevin in *C. elegans* chemosensory cilia

**DOI:** 10.1101/531848

**Authors:** Jona Mijalkovic, Erwin J.G. Peterman

## Abstract

Cytoplasmic dyneins play a role in a myriad of cellular processes, such as retrograde intracellular transport and cell division. Small-molecule cytoplasmic dynein antagonists, ciliobrevins, have recently been developed as tools to acutely probe cytoplasmic dynein function. Although widely used to investigate cytoplasmic dynein 1, far fewer studies explore the effect of ciliobrevin on cytoplasmic dynein 2 or IFT dynein. Here, we use ciliobrevin A to partially disrupt IFT dynein in the chemosensory cilia of living *C. elegans*. Acute, low-concentration ciliobrevin treatment results in shortening of cilia and reduction of transport velocity in both directions. After longer exposure to ciliobrevin, we find concentration-dependent motor accumulations and axonemal deformations. We propose that maintenance of ciliary length requires a high fraction of active IFT-dynein motors, while structural integrity can be preserved by only a few active motors.

## Introduction

Cytoplasmic dyneins are large (~1.4MDa), multi-subunit, microtubule minus-end directed ATP-driven motor proteins with a wide range of vital, cellular functions [1]. Cytoplasmic dynein 1 plays, for example, a role in the axonal transport of cargo (*e.g.* membranous organelles, mRNA and proteins) and cellular positioning of the mitotic spindle, chromosomes and centrosomes. A second type of cytoplasmic dynein, called cytoplasmic dynein 2 or IFT dynein, drives retrograde intraflagellar transport (IFT) in cilia and flagella [2], cellular structures that act as antennae to detect and respond to changes in the extracellular environment and require motor-driven IFT for their assembly and maintenance [3-5].

Recent advances in understanding cytoplasmic dynein function *in vivo* have largely relied on gene modification tools such as MosSCI, CRISPR and expression regulation using RNAi [6-9]. An important limitation of these methods is that they induce long-term cellular changes and thus cannot be used to probe the effect of acute and controlled (partial) dynein loss of function. To overcome this, small-molecule cytoplasmic dynein antagonists have been developed, enabling real-time motor inhibition [10-12]. Ciliobrevins are dihydroquinazolinone compounds discovered in a high-throughput screen for Hedgehog-signaling (Hh) pathway inhibitors [13]. In this screen, mouse fibroblast Hh-responsive cells treated with HPI-4 (later renamed ciliobrevin A) displayed shorter or absent cilia and accumulated transcription factor Gli2 and IFT-B particle subunit IFT88, pointing to perturbed retrograde transport [10, 13]. Ciliobrevin A and its derivative ciliobrevin D have since been utilized to investigate dynein-1-driven processes in cultured cells, such as chromosome segregation [14, 15], axonal transport and elongation [16] and centrosome orientation [17, 18]. Ciliobrevin was additionally found to inhibit IFT-mediated gliding motility and membrane-protein transport in *Chlamydomonas*, highlighting its potential as a dynein-2 inhibitor *in vivo* [19, 20]. However, the use of ciliobrevins in other living organisms to probe IFT-dynein function has remained underexplored.

Here, we test the efficacy of ciliobrevin A as IFT-dynein inhibitor in living *C. elegans* and examine the effect of acute dynein perturbation on IFT-motor ensemble distributions and velocities. We find that IFT in *C. elegans* phasmid chemosensory cilia is not halted by acute, low-concentration ciliobrevin treatment, but slows down, resulting in shortening of the ciliary axoneme. After prolonged ciliobrevin exposure, we find concentration-dependent motor accumulations and axonemal deformations indicative of more severely impaired transport. Based on these findings, we propose that maintenance of maximum ciliary length requires a high fraction of active IFT-dynein motors, while structural integrity can be preserved by only a few active motors.

## Results

### Optimization of ciliobrevin A dosing in *C. elegans*

First, we sought to determine the dose required to exert an inhibitory effect on IFT dynein in *C. elegans* phasmid cilia. Nematodes expressing fluorescently-tagged IFT-dynein light intermediate chain XBX-1 (XBX1::GFP) were initially treated with 20-100 µM ciliobrevin A, the concentration range reported to give maximum inhibitory response in different cell lines [10, 13, 16]. The animals were incubated in a ciliobrevin-containing solution or ciliobrevin-coated Nematode Growth Medium (NGM) plates with OP50 *E. coli* for 1 hour and then imaged using epifluorescence microscopy [21, 22]. We subsequently screened for changes in phasmid cilium length using long-exposure fluorescence images of XBX-1::EGFP. Visual inspection of the fluorescence images revealed that, in contrast to the cell lines studied before, no ciliary shortening was observed at these concentrations in *C. elegans*, suggesting that ciliobrevin does not elicit an inhibitory effect in this concentration range in the nematode. Exposure to increasingly higher doses resulted in observable ciliary shortening at concentrations above 0.7 mM, ~3-7 times higher than typically used in cell lines [10, 13, 16]. This higher concentration could suggest that ciliobrevin has a lower affinity for *C. elegans* IFT dynein, but it more likely reflects limited penetration of the drug or absorption-related drug loss in the nematode. Moreover, it is consistent with dosing regimens of other small-molecule drugs in *C. elegans* that have been reported to be in the millimolar-range [23, 24].

### Acute ciliobrevin treatment results in roadblocks, ciliary shortening and impaired bidirectional transport

To examine the effect of minimal but acute IFT-dynein inhibition, nematodes were subjected to 5-minute treatment with a “low”, 0.7 mM ciliobrevin dose (Fig 1A). Ciliobrevin is soluble in DMSO, a solvent that is toxic to *C. elegans* at concentrations above 2% V/V [25]. To test whether effects were specific to ciliobrevin and not induced by the 1% V/V DMSO used in the ciliobrevin experiments, we also treated nematodes with a control solution containing the same amount of DMSO. After dosing, the nematodes were allowed to recover briefly on unseeded NGM plates (2-3 minutes) and then anesthetized for 10 minutes using levamisole (Fig 1A). Ciliobrevin-induced ciliary shortening was quantified using long-exposure fluorescence images of XBX-1::EGFP (Fig 1B). In untreated nematodes, XBX1::EGFP is localized specifically to the chemosensory cilia and is distributed across the entire cilium [21, 22]. In control-treated nematodes, the average length occupied by XBX-1 was 8.33 ± 0.34 µm (average ± s.e.m), in agreement with previous length determination of *C. elegans* cilia [22, 26]. After acute, low-concentration ciliobrevin A treatment, the average length occupied markedly reduced to 5.85 ± 0.44 µm (Fig 1C). Our results confirm that, similar to observations in *Chlamydomonas* and ciliated cells, ciliobrevin can induce ciliary shortening in *C. elegans* [19].

**Fig 1.**
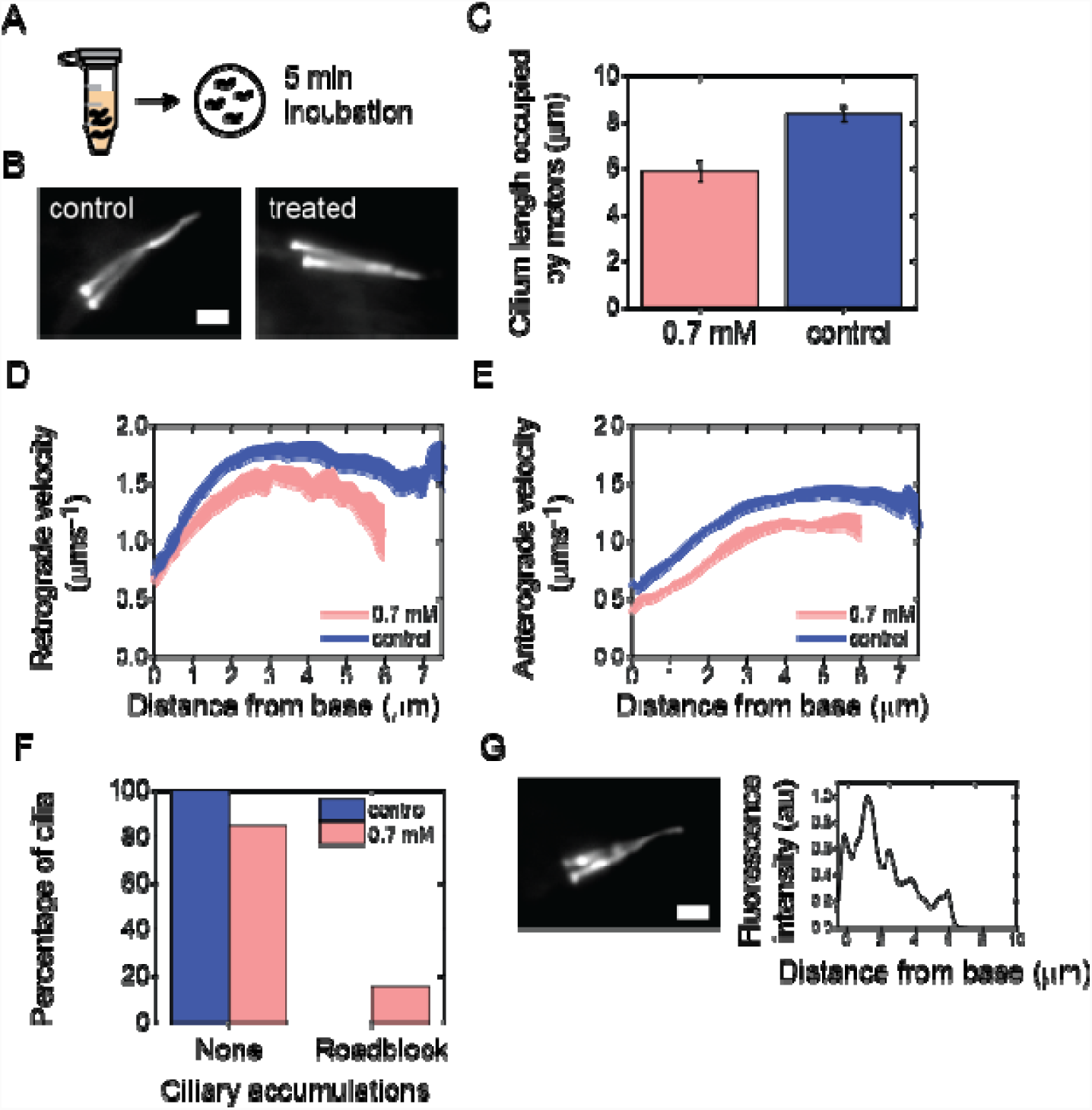
Acute ciliobrevin treatment results in roadblocks and ciliary shortening. **(A)** Dosing schematic: 5-minute exposure to 0.7 mM ciliobrevin A or control (2% DMSO in M9). **(B)** Example of summed fluorescence intensity images (150 subsequent images) of XBX-1 (XBX-1::EGFP) in cilia of treated and control animals. Scale bar 2 µm. **(C**) Average length occupied by XBX-1 after exposure to 70 mM ciliobrevin A (light red; n=11 cilia) or control (blue; n=4 cilia). Error is s.e.m. **(D)** Extent of XBX-1 accumulation in worms treated with 70 mM ciliobrevin A (light red) and control (2% (V/V) DMSO in M9; blue). **(D)** Average retrograde XBX-1 velocity after exposure to 70 mM ciliobrevin A (n=38 trains from 11 cilia; 8 worms) and control (n=24 trains from 4 cilia; 4 worms). **(E)** Average anterograde XBX-1 velocity after exposure to 70 mM ciliobrevin A (n=44 trains from 11 cilia; 8 worms) and control (n=24 trains from 4 cilia; 4 worms). Line thickness represents s.e.m. **(F)** Example summed fluorescence intensity image of XBX-1 (XBX-1::EGFP) and corresponding normalized fluorescence intensity profile showing roadblock accumulations. Scale bar 2 µm.

To determine whether the velocity of IFT trains is affected by acute ciliobrevin treatment, we next generated direction-filtered kymographs and extracted position-dependent velocities of XBX-1::EGFP using KymographClear and KymographDirect (Fig 1D, E) [27]. In the retrograde direction (tip to base, Fig 1D) IFT dynein is the active driver of transport, but in the anterograde direction (base to tip, Fig 1E) IFT dynein is carried as cargo by the kinesin-2 family motors OSM-3 and kinesin-II. We found that retrograde IFT-train velocity is reduced by ~12%, suggesting rapid inhibition of (at least some) IFT-dynein motors participating in transport. Interestingly, kinesin-2-driven anterograde transport was also reduced (~21%). While the original report by Firestone *et al* suggested that ciliobrevin A and D do not affect kinesin-1-dependent microtubule gliding in *in vitro* assays with only kinesin-1, our data is consistent with studies in primary neurons showing that ciliobrevin D affects both anterograde and retrograde transport [16, 28]. Similarly, in *Chlamydomonas*, anterograde transport impairment occurs already 2 minutes after treatment [19]. It is possible that ciliobrevin is not specific to the cytoplasmic dynein ATP binding site, but interpretation of these findings is difficult owing to the strong interdependence between anterograde and retrograde transport [29]. Functional IFT requires continuous turnover and impairment of retrograde transport could thus indirectly lead to impairment of anterograde transport [30]. Another possibility is that anterograde IFT trains require functional, uninhibited dynein motors [31].

Next, we used the long-exposure fluorescence images to visualize the effect of ciliobrevin A on XBX-1 distribution. In all control-treated cilia there were no visible accumulations of motors along the cilium (Fig 1B, F). In 15.4% of the acutely ciliobrevin-treated cilia, however, we observed XBX-1 accumulations at non-specific locations along the cilium (Fig 1F, G). We attribute these “roadblock” accumulations to inhibition of a small subset of IFT-dynein motors, resulting in stalled IFT trains. These non-moving IFT trains can be bypassed by active motors (Movie S1), indicating that they do not completely block IFT. Taken together, our findings affirm the efficacy of ciliobrevin A in the phasmid cilia of living *C. elegans*. Importantly, we show that acute IFT-dynein inhibition using a minimal ciliobrevin concentration does not completely halt IFT. While the number of functional dynein motors is likely diminished, IFT is maintained, albeit at a lower velocity, resulting in a reduced ciliary length.

### Prolonged ciliobrevin A treatment leads to concentration- and velocity-dependent ciliary shortening

To explore the effect of more prolonged inhibition of IFT dynein, we subjected nematodes to 1 hour treatment with low- and high-concentration ciliobrevin (0.7 and 1.4 mM in 2% DMSO, respectively), or 2% DMSO solution as control. Due to the limited solubility of ciliobrevin and DMSO toxicity (above 2% V/V), 1.4 mM was the highest ciliobrevin concentration attainable in our experiments. In control-treated nematodes, the average distance occupied by motors was 8.25 ± 0.18 µm, similar to control experiments with 1% DMSO (Fig 2A). After 1-hour treatment with 0.7 mM and 1.4 mM ciliobrevin, cilia appeared shorter and the average length occupied by XBX-1::EGFP reduced to 6.91 ± 0.17 µm and 6.11 ± 0.30 µm respectively. To probe whether the observed shortening is due to motor retraction or axonemal shortening, we also exposed animals with labeled ciliary tubulin TBB-4::EGFP to 1.4 mM ciliobrevin solution. The average cilium length in these animals was 5.59 ± 0.22 µm (Fig 2A), confirming that ciliobrevin induces retraction of the axoneme. To obtain further insight into ciliary shortening, we plotted the distribution of post-treatment length occupied by XBX-1 for each treatment (Fig 2A). While in all low-concentration-treated worms XBX-1 extended from base to beyond the proximal segment, in some high-concentration treated worms we observed XBX-1 only occupying the first 2 – 4 µm of the cilium, most likely indicating retraction of (part of) the proximal segment. Such severe truncations were not observed with acute or prolonged low-concentration treatment, revealing concentration-dependent ciliary shortening in response to prolonged ciliobrevin treatment (Fig 1C, Fig 2A).

**Fig 2.**
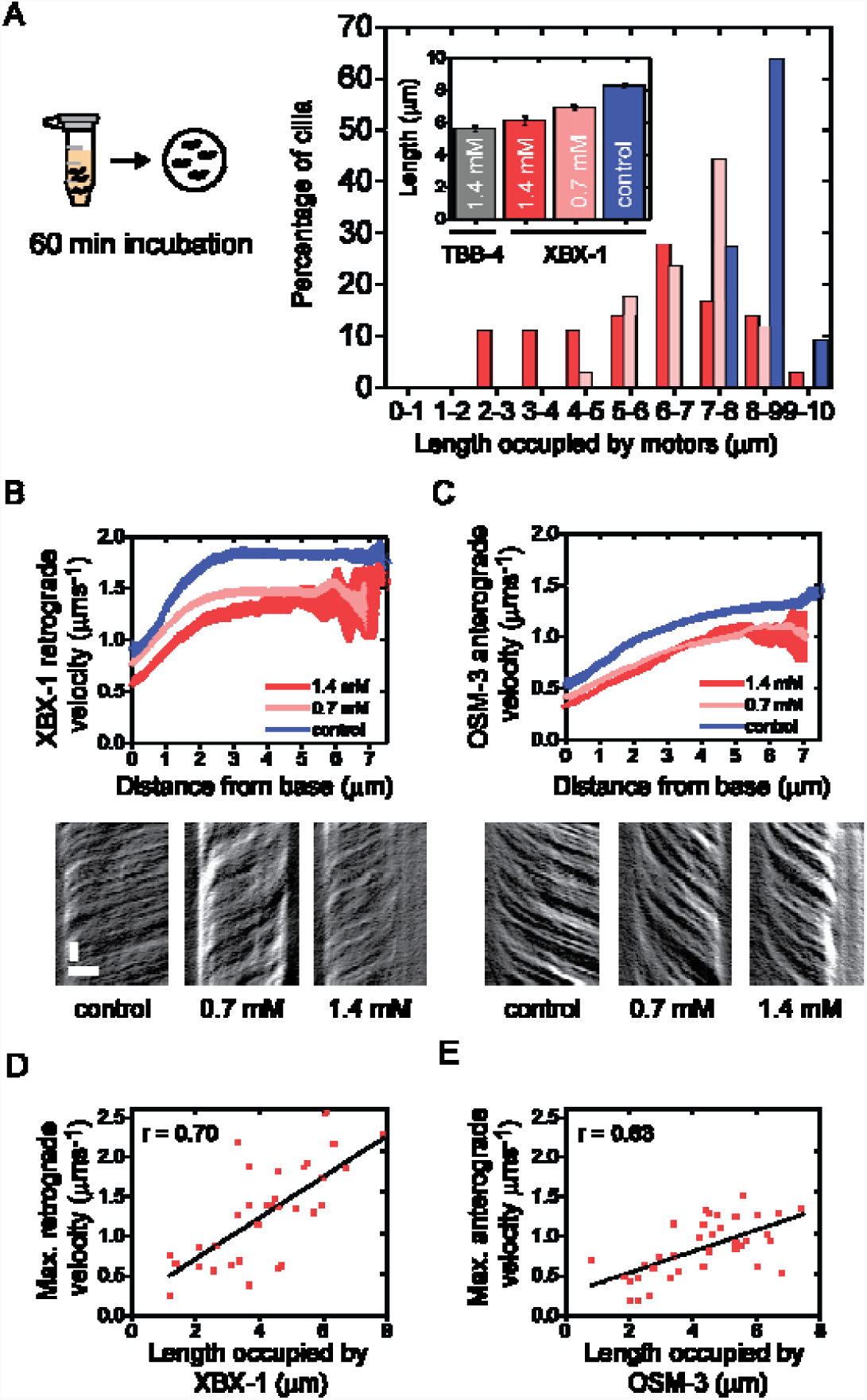
Concentration- and velocity-dependent ciliary shortening after ciliobrevin A exposure. **(A)** Dosing schematic: 60-minute exposure to ciliobrevin A or control (2% DMSO in M9). Inset: Average cilium length (TBB-4, grey, n=39) and average ciliary length occupied by XBX-1 after 60-minute exposure to 0.7 mM ciliobrevin A (light red, n=34), 1.4 mM ciliobrevin (red, n=39) or control (blue). Error is s.e.m. Graph: Distribution of cilium lengths occupied by XBX-1 after 60-minute exposure to ciliobrevin A or control. **(B)** Train-averaged XBX-1 retrograde velocity after exposure to 0.7 mM ciliobrevin A (n=130 trains from 34 cilia; 18 worms), 1.4 mM ciliobrevin A (n=138 trains from 35 cilia; 19 worms) and control (n=79 trains from 11 cilia; 7 worms). Line thickness represents s.e.m. Representative corresponding kymographs are shown. Time: vertical; scale bar 2 s. Position: horizontal; scale bar 2 µm. **(C)** Train-averaged OSM-3 anterograde velocity and representative kymographs after exposure to 0.7 mM ciliobrevin A (n=136 trains from 34 cilia; 18 worms), 1.4 mM ciliobrevin A (n=146 trains from 35 cilia; 19 worms) and control (n=44 trains from 11 cilia; 7 worms). Line thickness represents s.e.m. **(D-E)** Maximum XBX-1 (D) and OSM-3 (E) velocity versus ciliary length occupied by motors. Red squares: data points; black lines represent regression lines obtained from a Pearson’s correlation analysis (correlation coefficient r: 0.70 for XBX-1 and 0.63 for OSM-3).

We next investigated the effect of prolonged ciliobrevin treatment on IFT-motor velocity using a dual-label *C. elegans* strain with fluorescently-tagged IFT dynein and one of the anterograde kinesin-2 motors, OSM-3 [22]. We recorded fluorescence image sequences of IFT dynein (XBX-1::EGFP) and OSM-3 (OSM-3::mCherry) and generated kymographs. Position-dependent velocities were then extracted using KymographClear and KymographDirect (Fig 2B, C) [27]. Prolonged treatment with ciliobrevin resulted in a ~24% (at 0.7 mM) and ~36% (at 1.4 mM) lower retrograde IFT-dynein velocity (Fig 2B). The anterograde (OSM-3) velocity was also reduced, by ~31% (at 0.7 mM) and ~34% (at 1.4 mM; Fig 2C). These results show that higher concentrations of ciliobrevin and longer exposure to the drug result in a larger reduction of the velocity in both directions. To further explore the relationship between cilium length and IFT velocity, we plotted the maximum anterograde (Fig 2D) and retrograde (Fig 2E) velocity for each cilium. We observe a clear trend (Pearson’s r = 0.70 and 0.63 for XBX-1 and OSM-3 respectively) between maximally attained IFT velocities and length of cilium occupied by motors, suggesting that the extent of ciliary shortening could be velocity-dependent.

### Prolonged ciliobrevin A exposure leads to altered motor distribution

To investigate the effect of prolonged ciliobrevin A exposure on IFT-dynein distribution, we characterized the extent of IFT-dynein accumulations after high- and low-concentration treatment (Fig 3). As expected, in control-treated nematodes there were no aberrant motor accumulations (2% DMSO; Fig 3A). In nematodes that underwent prolonged, low-concentration treatment, 50% of phasmid cilia had no motor accumulations and 29% had “roadblock” accumulations at apparently random positions along the cilium. In a smaller fraction of the cilia (9%) we observed IFT-dynein accumulations only at the ciliary tip (Fig 3B), consistent with defective retrograde transport. The observation of tip accumulation of IFT components upon ciliobrevin treatment is consistent with earlier studies in cell lines[10] and *Chlamydomonas* [19]. In some cilia (12%), motor accumulation in the distal segment was more substantial, covering at least 1 µm of the distal segment including tip (Fig 3C). In nematodes that underwent prolonged, high-concentration ciliobrevin treatment we observed more severe disruptions of the IFT-dynein distribution. 13% of the cilia showed motor accumulation only at the tip, while in 33% of cilia dynein motors had stalled in the entire distal segment, substantially affecting the integrity of the axoneme (Fig 3C). We hypothesize that in these nematodes there are too few active IFT-dynein motors left to maintain bidirectional IFT, resulting in (partial) collapse of the axoneme. In some cases, we observed a remarkable heterogeneity in the response to ciliobrevin between the two cilia in a phasmid pair within one organism, despite identical exposure. In the example shown in Fig 3B, one phasmid cilium is substantially shortened while the other has maintained its full length. These observations suggest that there is a balance point in the number of functional IFT components required for distal segment maintenance and structural integrity (Fig 3F). In this view, only a small fluctuation of component numbers below this balance point, in a given cilium at a given time, will result in collapse of IFT and ciliary structure. In addition, after prolonged high-concentration treatment (but not in acute or prolonged low-concentration treatment) we observed structural aberrations of the cilia in some nematodes, such as tight bends (Fig 3E). This could be due to a ciliobrevin-induced effect on tubulin structure or disruption of IFT-mediated tubulin transport [16, 32] at these high concentrations.

**Fig 3.**
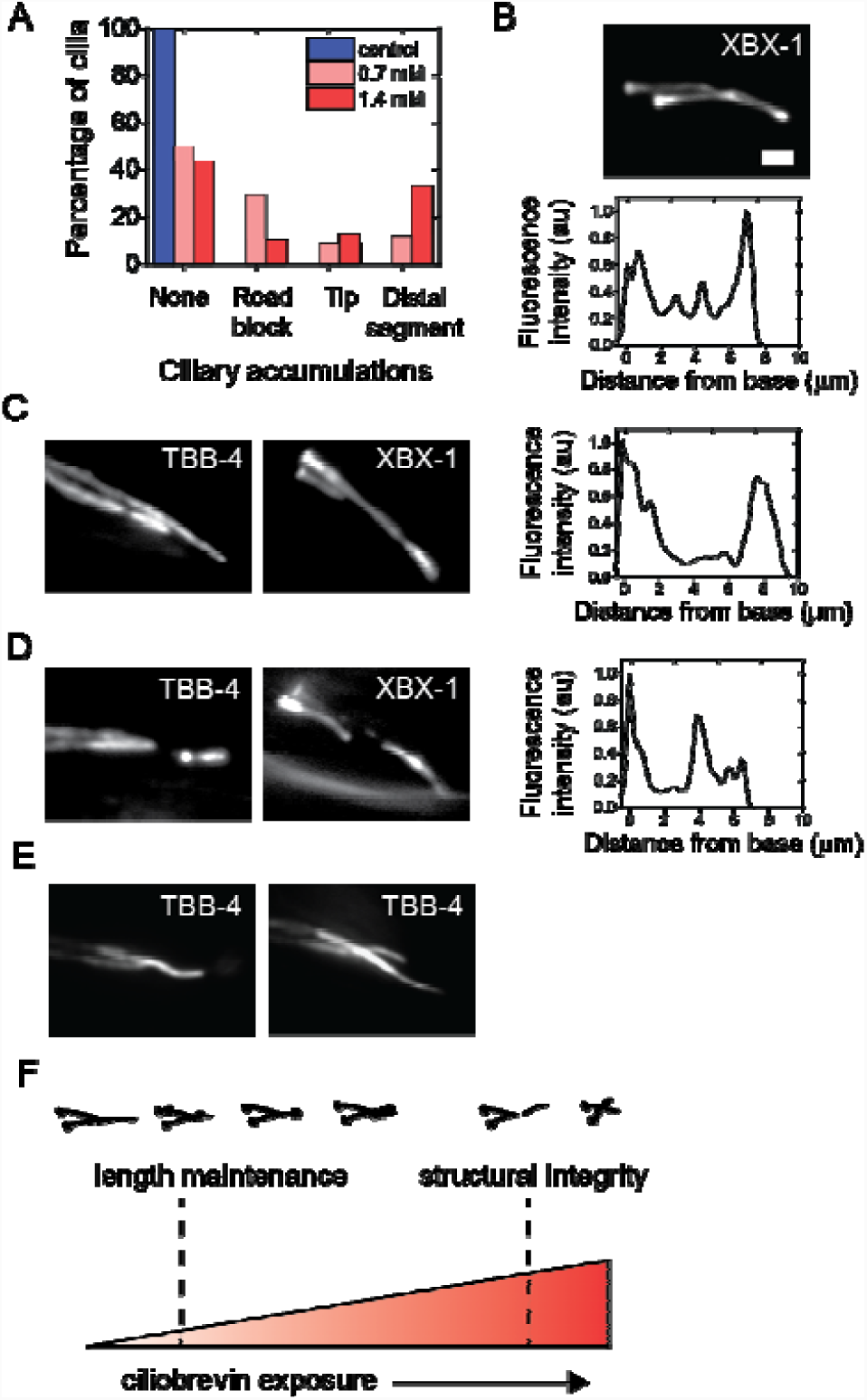
Concentration-dependent effects on motor distribution and the ciliary axoneme. **(A)** Extent of XBX-1 accumulation in worms treated with 0.7 mM ciliobrevin A (light red), 1.4 mM ciliobrevin A (red) and control (2% DMSO in M9; blue). **(B-D)** Example summed fluorescence intensity images of XBX-1 (XBX-1::EGFP) and TBB-4 (TBB-4::EGFP) and corresponding normalized XBX-1 fluorescence intensity profile showing tip accumulation **(B)** and distal segment accumulation **(C, D). (E)** Example summed fluorescence intensity images of TBB-4 after 1.4 mM ciliobrevin treatment showing axonemal malformations. Scale bar 2 µm. **(F)** Model of ciliobrevin-induced IFT-dynein inhibition on IFT. The axonemal length maintenance balance-point is easily disturbed, whereas structural integrity can be maintained with a much lower number of active IFT-dynein motors.

Taken together, our findings enable us to propose a model of the ciliary effects of IFT-dynein inhibition by ciliobrevin. Acute, low-concentration treatment results in axonemal shortening coupled with a reduction in IFT velocity in both directions, suggesting that both maximum ciliary length and IFT-velocity maintenance require a relatively high number of active IFT-dynein motors. At higher ciliobrevin concentrations, motors become slower and cilia become even shorter, until only a few active motors remain, tipping the balance of ciliary structure, resulting in severely truncated cilia with axonemal defects.

## Discussion

The recent development of ciliobrevins has made it possible to acutely inhibit cytoplasmic dynein in several cellular systems [10]. Here, we show that ciliobrevin A, dosed in the low millimolar range, can exert an inhibitory effect on IFT dynein in *C. elegans* phasmid chemosensory cilia.

Although we have shown that ciliobrevin A can inhibit IFT dynein in living *C. elegans*, there are several caveats to using it as a quantitative and well-controllable tool to probe dynein function. Firstly, the pharmacokinetics of ciliobrevin in *C. elegans* is unknown and ciliobrevin might not be specific to one cytoplasmic dynein subtype. Therefore, if uptake primarily takes place through the mouth instead of the ciliary openings, dynein-1-driven function and dendritic transport could also be affected. Additionally, the absolute intracellular ciliobrevin concentration is unknown and could be up to 40 times lower than the extracellular dose [24]. Secondly, in IFT, motors work in teams of several tens of motors and we cannot currently determine the number of motors actively engaged in transport. As such, ciliobrevin concentration is the only surrogate for the number of (in)active motors, but this relationship may not necessarily be linear.

Notwithstanding these limitations, we have made some clear observations of the effects of ciliobrevin on IFT in *C. elegans* phasmid cilia. The first effects of acute ciliobrevin treatment are that IFT velocity has decreased 12-21%, both in anterograde and retrograde direction, and that cilia are ~30% shorter. These effects occur before IFT motors accumulate at the ciliary tip, as we observe after longer exposure, consistent with previous studies on cells [10]. As a consequence, these first effects represent the initial response of the IFT system to minimal IFT-dynein perturbation. A lower IFT-dynein velocity can be attributed to frictional drag or less efficient collective motion due to ciliobrevin-induced inhibition of a subset of motors [33]. The lower velocity appears to correlate with the retraction of the axoneme. This suggests that a minimum IFT-train velocity (and potentially a minimum number of active IFT-dynein motors) is required for the maintenance of the full length of the cilium. Reduced IFT velocities have also been observed in *Chlamydomonas* mutants with shorter flagella[34]. In *Chlamydomonas* experiments where flagellar shortening was induced by a pH shock, long flagella also had higher IFT velocities than shorter flagella [35]. IFT trains, however, were larger in short than in long flagella, leading to a balance-point model based on train size [35]. To connect our findings with those in *Chlamydomonas*, it would be interesting to detect how many motors are actively engaged with the axoneme at a given time on an IFT train, but such tools are currently lacking *in vivo*.

Our results suggest that IFT is relatively robust. Despite inhibition of a subset of IFT-dynein motors, bidirectional transport still continues, albeit at a lower velocity, resulting in a shorter axoneme. Such adaptations to mild perturbations of ciliary processes could be necessary for quick responses to extracellular changes.

## Materials and Methods

### *C. elegans* strains and dosing

*C. elegans* were maintained according to standard procedures. Nematodes were grown at 20°C on Nematode Growth Medium (NGM) plates seeded with *Escherichia coli* OP50 bacteria. The strains used in this study, EJP212 (XBX-1::EGFP, OSM-3::mCherry) [22] and EJP401 (TBB4::EGFP), were generated using Mos1-mediated single-copy insertion (MosSCI) [36]. Ciliobrevin A powder (H4541, Sigma Aldrich) was dissolved in fresh DMSO at maximum solubility to make a 140 mM stock solution stored at −20°C for a maximum of 1 month (after 1 month we observed loss of efficacy). The ciliobrevin dosing solution was made by diluting this stock solution in M9 to 1.4 mM (2% (V/V) DMSO content) or 0.7 mM (1% DMSO content) and used the same day. Control (vehicle) dosing solutions were made by dissolving DMSO in M9 (1% and 2% DMSO). Young adult hermaphrodites were dosed by transferring to standard 0.2 ml PCR tubes (Thermo Scientific) with 0.7 mM or 1.4 mM ciliobrevin solution for 5 min or 60 min. Nematodes were allowed to recover on unseeded NGM plates for 2-3 minutes post-dosing.

### Fluorescence imaging and analysis

*C. elegans* were anesthetized with 5mM levamisole in M9 and immobilized on a 2% agarose in M9 pad covered with a 22 × 22□mm cover glass and sealed with VaLaP. Fluorescence imaging was done using a custom-built epi-illuminated fluorescence microscope as described previously [22, 26]. Fluorescence images were analyzed using KymographDirect and KymographClear [27].

## Acknowledgements

We acknowledge financial support from the Netherlands Organization for Scientific Research (NOW) via a Vici grant (E.J.G.P.). We thank Pierre Mangeol for help with the dosing set-up in the initial stages of the project.

